# Pervasive divergence in protein thermostability is mediated by both structural changes and cellular environments

**DOI:** 10.1101/2024.12.09.627561

**Authors:** Nilima Walunjkar, Timothy Y. Lai, Nasima Akhter, James H. Miller, John Q. Bettinger, Erin Marcus, Eric M. Phizicky, Sina Ghaemmaghami, Justin C. Fay

**Affiliations:** Department of Biology, University of Rochester, Rochester, NY 14610; Department of Biochemistry and Biophysics, Center for RNA Biology, University of Rochester School of Medicine, Rochester, NY 14610

## Abstract

Temperature is a universal environmental constraint and organisms have evolved diverse mechanisms of thermotolerance. A central feature of thermophiles relative to mesophiles is a universal shift in protein stability, implying that it is a major constituent of thermotolerance. However, organisms have also evolved extensive buffering systems, such as those that disaggregate and refold denatured proteins and enable survival of heat shock. Here, we show that both cellular and protein structural changes contribute to divergence in protein thermostability between two closely related *Saccharomyces* species that differ by 8°C in their thermotolerance. Using thermal proteomic profiling we find that 85% of *S. cerevisiae* proteins are more stable than their *S. uvarum* homologs and there is an average shift of 1.6°C in temperature induced protein aggregation. In an interspecific hybrid of the two species, *S. cerevisiae* proteins retain their thermostability, while the thermostability of their *S. uvarum* homologs is enhanced, indicating that cellular context contributes to protein stability differences. By purifying orthologous proteins we show that amino acid substitutions underlie melting temperature differences for two proteins, Guk1 and Aha1. Amino acid substitutions are also computationally predicted to contribute to stability differences for most of the proteome. Our results imply that coordinated changes in protein thermostability impose a significant constraint on the time scales over which thermotolerance can evolve.

## Introduction

Microorganisms vary widely in the temperatures they experience and their thermal limits. Protein stability is a key component of thermotolerance as heat causes protein unfolding, aggregation and loss of fitness. Consequently, there are diverse mechanisms of achieving protein homeostasis at different temperatures. For example, the heat shock response is a near universal response to temperature-induced protein denaturation and is tuned to an organism’s upper thermal limit [1–5]. Additionally, the proteomes of thermophiles are exceptionally robust to temperature induced aggregation [6–8]. Given the frequent and sometimes rapid temperature shifts due to climate change, evolution of protein stability is an important but complex variable relevant to an organism’s thermal tolerance.

Protein stability is influenced by both protein structure and the cellular environment. Most amino acid substitutions destabilize proteins or make it more difficult for them to fold into their native conformations [9–11]. However, both natural and experimental evolution show that proteins can attain higher thermostability without compromising their activity [12,13]. The cellular environment can impact protein stability through pH [14], ligand or protein interactions [15,16], post-translational modifications [17], thermoprotectant molecules [18], and protein chaperones [19]. Because most proteins are marginally stable due to the small changes in free energy (ΔG) between folded and unfolded states [20], both structural changes and cellular factors may continuously evolve with the temperatures experienced by an organism [21,22].

The numerous ways in which stability can be modulated provide multiple evolutionary paths to thermal adaptation. Cellular chaperones and thermoprotectants can yield immediate stabilization of proteins but may be limited to small, temporary or partial stabilization of the proteome [23,24]. Structural stabilization of proteins can be broadly achieved through amino acid substitutions, but may be limited by the time and coordination of changes across the proteome. While thermal adaptations have been documented for both paths [25–28], experimental evolution in multiple species has resulted in limited increases in upper thermal limits [28–34]. This indicates that protein stability might constrain the rate of temperature adaptation and raises the possibility that proteome-wide stabilization may be required for natural systems to attain systemic increases in thermotolerance.

A major challenge to understanding divergence in protein stability between species is disentangling cellular contributions to stability from those that are structurally encoded by a protein’s amino acid sequence. Although changes in stability have been characterized through purification of individual proteins [25,35], proteomic measures of melting temperature divergence between thermophiles and mesophiles do not always distinguish between structural and cellular contributions. In addition to chaperones and thermoprotectants the cell environment can influence protein stability through post-translational modifications [17], the presence of binding partners [15], macromolecular crowding or other constituents of the cell. Proteomic studies have also shown that protein melting temperatures are somewhat dependent on the cell type and environment [15,36,37], and that melting temperatures of proteins from mesophilic species are not closely tied to their optimal growth temperature [6,38]. This has made it difficult to understand the extent to which protein stability co-evolves with an organism’s thermal niche.

In this study we characterize proteome stabilities of two thermally diverged *Saccharomyces* species, *S. cerevisiae* and *S. uvarum*. These two species have diverged for 16 million years [39] and have evolved an 8°C difference in their thermal growth limits (Figure 1), resulting in substantial temperature-dependent fitness differences [40] and their differential use in low versus higher temperature fermentations [41,42]. The two species can also produce viable hybrids, enabling protein stability to be measured in the same cellular environment. Our results provide direct evidence that both structural and cellular changes contribute to widespread changes in protein thermal stability in yeast and imply that protein stability is likely a major constraint on adaptation to high temperatures.

**Figure 1:**
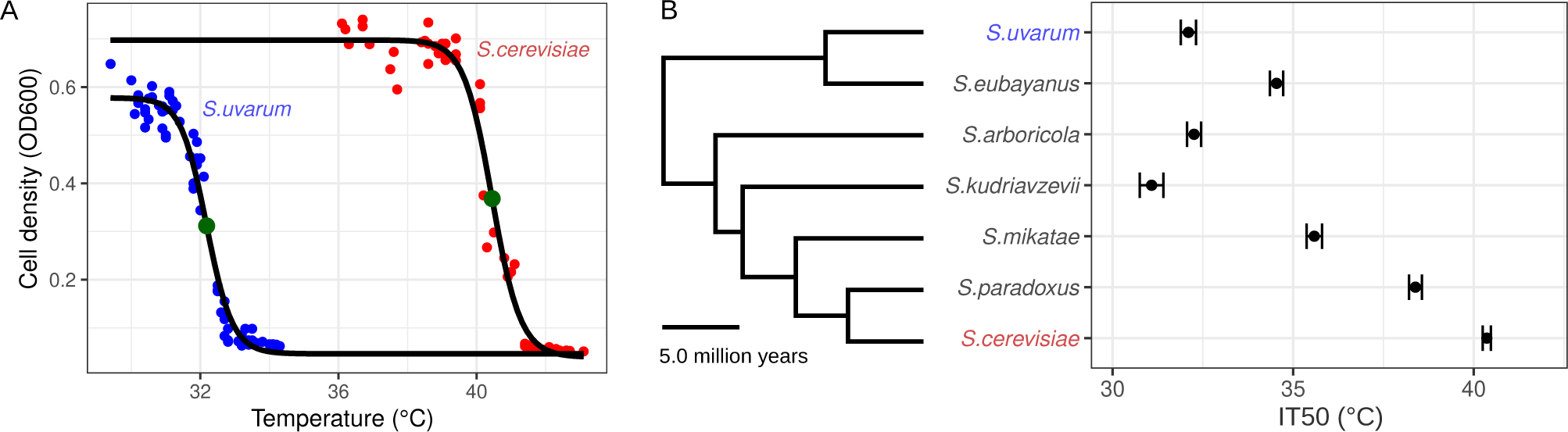
Divergence in thermotolerance among the *Saccharomyces* species. (A) Thermotolerance was measured by the temperature at which growth yield (OD600) was reduced by 50% (IT50, green dots), based on treatments using a gradient thermocycler. (B) *Saccharomyces* species IT50 with whiskers showing 95% confidence intervals. Species tree and divergence times were obtained from Shen et al. [39].

## Results

### *S. cerevisiae* proteins are more thermostable than their *S. uvarum* orthologs

To investigate divergence in protein stability between *S. cerevisiae* and *S. uvarum* we used thermal proteomic profiling to measure protein melting temperatures [16]. Cell extracts were subjected to temperature treatments and the soluble fraction of proteins was quantified by LC-MS/MS. We estimated the temperature at which the levels of each protein in the soluble fraction was reduced by half and obtained melting temperatures (Tm) for 1,330 (replicate 1) and 914 (replicate 2) orthologous protein pairs (Table S1).

On average, *S. cerevisiae* proteins had a 1.6°C higher melting temperature, and 85% of *S. cerevisiae* proteins were more thermostable than their *S. uvarum* orthologs (Binomial p < 0.001, n = 827, Figure 2A, Table S2). For proteins with melting temperatures that had non-overlapping confidence intervals between species (n = 206), 90% of *S. cerevisiae* proteins were more stable than *S. uvarum* (Figure 2A, Table S2). The large fraction of proteins with higher melting temperatures in *S. cerevisiae* indicates uniform divergence in proteome stability. However, mitochondria and membrane-associated proteins were enriched for higher melting temperatures in *S. cerevisiae* whereas ribosomal proteins were enriched for higher melting temperatures in *S. uvarum* (Table S3). Larger melting temperature differences were associated with lower protein abundance and higher melting temperatures (Table S4).

**Figure 2:**
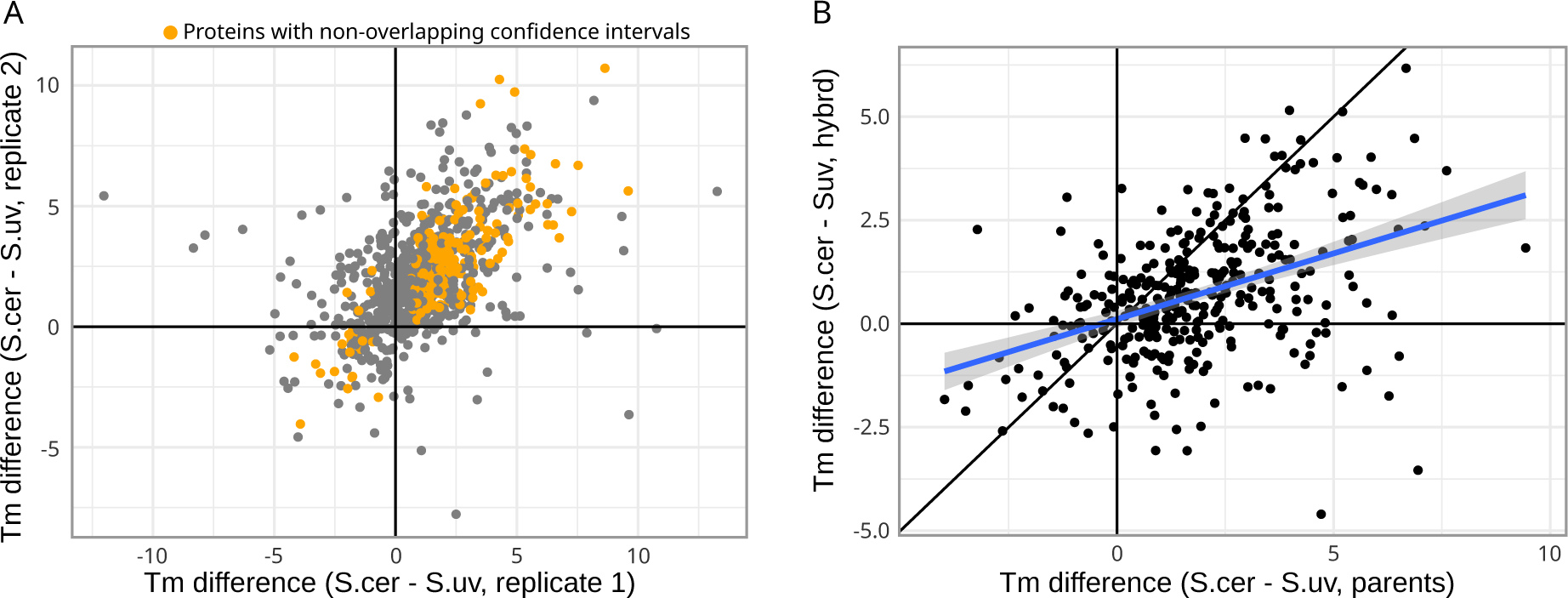
*S. cerevisiae* proteins have higher melting temperatures than *S. uvarum*. (A) Melting temperature (Tm) differences between *S. cerevisiae* and *S. uvarum* for replicate 1 versus replicate 2. Orange points show proteins with melting temperature confidence intervals that did not overlap between species in both replicates. (B) Hybrid Tm difference compared to the average of the two parental Tm differences. The blue line shows the linear regression (slope = 0.32).

Previously, *S. uvarum* hexokinases were shown to have lower thermal stability compared to *S. cerevisiae* but paralogous hexokinases weren’t distinguished [43]. In our proteomic data hexokinases (Hxk1, Hxk2, Glk1 and Emi2) from *S. cerevisiae* show higher stability than their *S. uvarum* orthologs (Table S5). As an additional assessment of stability differences we measured enzyme activities of malate dehydrogenase (encoded by *MDH1*, *MDH2* and *MDH3*) and glutathione reductase (*GLR1*) following heat treatment of cell extracts. Consistent with our proteomics data (Table S5), *S. cerevisiae* activity was more resistant to temperature than *S. uvarum* (Figure S1).

### Proteome stability depends on the cellular environment

Protein stabilities can depend on the cellular environment, due to post-translational modifications, such as glycosylation [17], or the stability of other proteins in the same complex [15]. To control for the cellular environment, we also measured the melting temperatures of proteins from an *S. cerevisiae* x *S. uvarum* hybrid and obtained species-specific measurements of orthologous protein pairs using species-specific peptides. In the hybrid, 67.4% of detected *S. cerevisiae* proteins were more stable than their *S. uvarum* orthologs (Table S2), and when combined with the parental data 87.7% of *S. cerevisiae* proteins were consistently more stable than their *S. uvarum* ortholog across both parental replicates and the hybrid (Binomial p < 2.2e-16, n = 204, Figure 2B). However, melting temperature differences between orthologs were smaller in the hybrid compared to their parents, indicating that other factors besides a proteins’ sequence contribute to the observed melting temperature differences between species.

We tested whether *S. cerevisiae* proteins were destabilized or *S. uvarum* proteins were stabilized in the interspecific hybrid. We found no consistent difference between the hybrid and parental stability of *S. cerevisiae* proteins, but *S. uvarum* proteins were significantly more stable in the hybrid compared with their parental cellular environment (Figure 3, Paired t-test, p < 2.2e-16). Given the stabilization of *S. uvarum* proteins in the hybrid, we tested whether hybrid stabilized proteins were associated with glycosylation, membranes, protein complexes, the ribosome, or the mitochondria. We found that only membrane proteins were more stable in the hybrid (Table S6), but this was true for both *S. uvarum* (OR = 4.31, Fisher’s exact test p = 0.0007) and *S. cerevisiae* proteins (OR = 2.78, Fisher’s exact test p = 0.0013). There was no significant difference in the membrane enrichment of *S. uvarum* compared to *S. cerevisiae* proteins (Logistic regression, p = 0.46).

**Figure 3:**
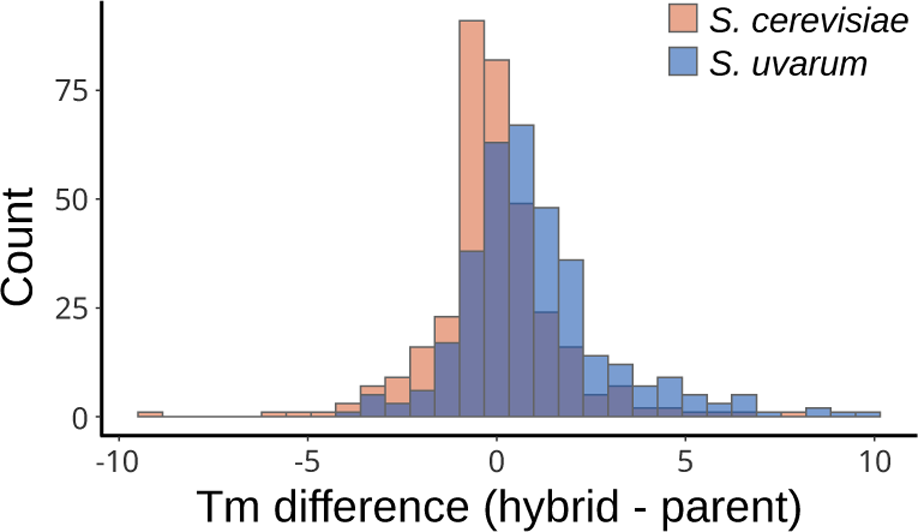
*S. uvarum* proteins are stabilized in the hybrid environment. The histogram shows the distribution of melting temperature differences between the hybrid and the parental species environments for both *S. cerevisiae* and *S. uvarum* proteins.

### Guk1 and Aha1 stability differences are structurally encoded

Many proteins exhibit consistent stability differences between the parents and between parental alleles within the hybrid, suggesting that they are structurally encoded by amino acid differences between species. To assess divergence in protein stability due to sequence alone, we selected two proteins (Guk1 and Aha1) and cloned and purified both from *S. cerevisiae* and *S. uvarum*. Guk1 and Aha1 were chosen as they are both small proteins with large and consistent melting temperature differences (Table S5). For each protein we used circular dichroism spectrometry to characterize their in-vitro melting curves by tracking ellipticity as a function of temperature. *S. cerevisiae* Guk1 had a 5.6°C higher melting temperature than that of *S. uvarum* (Figure 4A), demonstrating that the 21 amino acid substitutions have together resulted in divergence of protein stability. For Aha1, the *S. cerevisiae* protein was also more thermostable than that of *S. uvarum* due to 54 amino acid differences and an insertion/deletion (Figure 4B, Tm difference of 9.1°C). We also measured Guk1 activity following thermal treatments of purified protein and confirmed a loss of activity consistent with the protein’s melting temperature (Figure S1).

**Figure 4:**
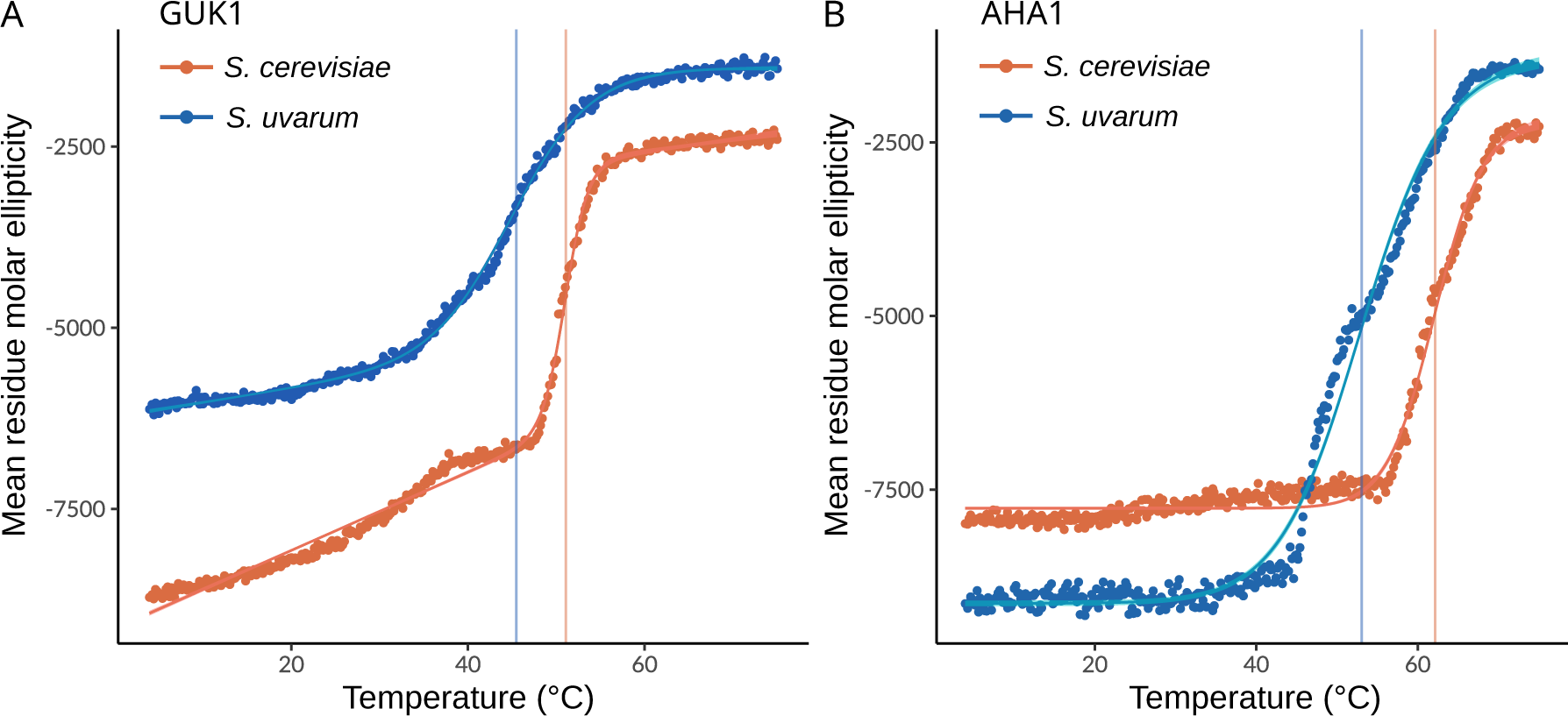
Divergence in Guk1 and Aha1 thermal stability. Circular dichroism was used to measure thermal loss of secondary structure for *S. cerevisiae* and *S. uvarum* Guk1 (A) and Aha1 (B). Secondary structure was measured by mean residue molar ellipticity (deg cm^2^ dmol^-1^ per residue) at 222 nm as a function of temperature (°C).

### Divergence in Guk1 protein thermostability does not affect fitness in *S. cerevisiae*

Guanylate kinase (Guk1) is an essential protein that catalyzes the conversion of guanosine monophosphate (GMP) to guanosine diphosphate (GDP). To test if Guk1 stability differences between *S. cerevisiae* and *S. uvarum* can cause temperature-dependent growth differences, we replaced the endogenous *S. cerevisiae GUK1* with the less stable *S. uvarum* allele under the *S. cerevisiae* promoter. There were no growth differences under various conditions and temperatures, or upon heat shock at 50°C (Figure S2). An allele-replacement in *S. uvarum* also showed no change in thermotolerance (Figure S3).

### Predictable differences in protein stability

If the higher melting temperatures of *S. cerevisiae* proteins are structurally encoded by amino acids differences between species, then replacement of *S. cerevisiae* amino acids with those present in *S. uvarum* should destabilize the protein. To assess the impact of amino acid substitutions on stability we generated AlphaFold2 structures for *S. cerevisiae* and *S. uvarum* proteins [44,45] and predicted the difference in ΔG (ΔΔG) using the structure based ACDC-NN method [46]. We also generated predictions for substitutions of *S. cerevisiae* amino acids into predicted *S. uvarum* protein structures to avoid any reference structure bias.

The average protein has 64 amino acid differences between the two species, corresponding to 87% identity, and 1.9 more stabilizing amino acids in *S. cerevisiae* compared to *S. uvarum* (Figure 5A). Most amino acid changes have small effect sizes (|ΔΔG| < 0.5), but the sum of the ΔΔG values for each protein are typically greater than 0.5 and show that 62.6% of *S. cerevisiae* proteins are predicted to be more stable than their *S. uvarum* orthologs (Figure 5B, Binomial p < 1e-15). Over the entire proteome, the cumulative differences in stability come from both small (| ΔΔG| < 0.5) and large effect changes (|ΔΔG| > 0.5, Figure 5C). These stability differences are mostly derived from regions with high confidence in AlphaFold structures and are also found in ACDC-NN-Seq ΔΔG predictions based on sequence alone (Table S7 and S8).

**Figure 5:**
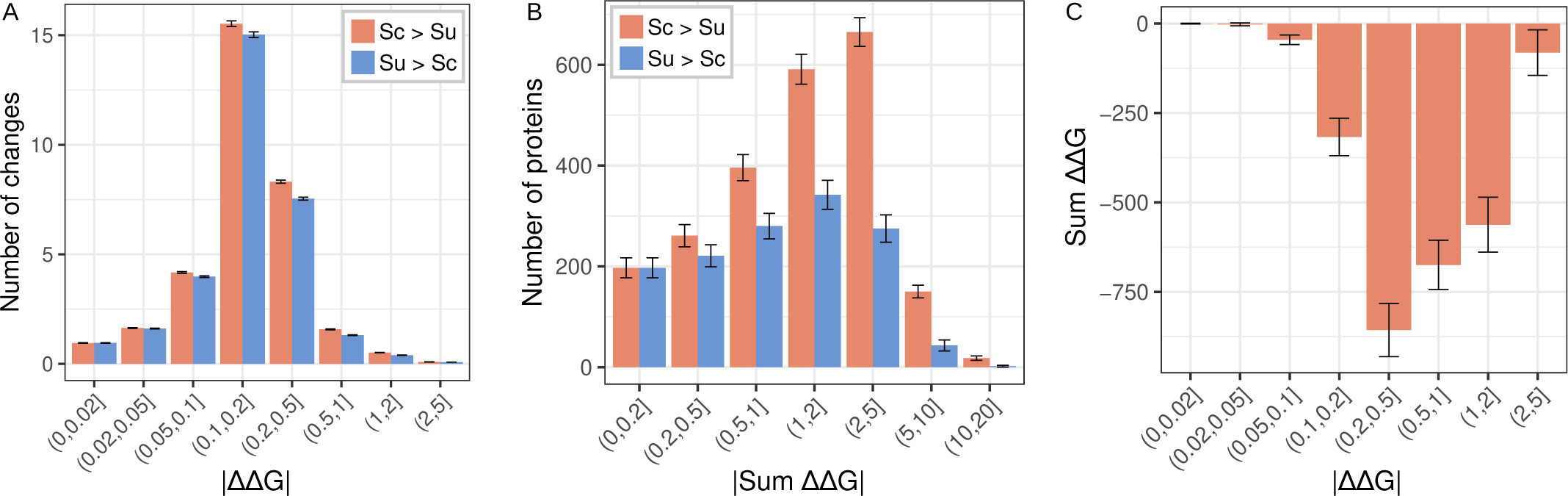
Distribution of protein stability differences between species. (A) Distribution of the average number of amino acid changes per protein where the *S. cerevisiae* allele is more stable than *S. uvarum* (Sc > Su) or vice versa (Su > Sc), binned by absolute value of ΔΔG. Whiskers show the 95% confidence interval based on the standard error. (B) Protein stability differences were measured by the sum of the ΔΔG values predicted by ACDC-NN for each amino acid change. The number of proteins with higher stability in *S. cerevisiae* or *S. uvarum* is shown grouped by the absolute value of ΔΔG. Whiskers show the binomial 95% confidence intervals. (C) Sum of positive ΔΔG and negative ΔΔG values, binned by absolute value of ΔΔG. Whiskers show bootstrap estimates of the standard error. A negative sum indicates *S. uvarum* alleles are cumulatively destabilizing.

Changes in ΔG can be correlated with changes in melting temperature depending on how the enthalpy and entropy of unfolding changes with temperature [47]. However, we found no correlation between the summed ΔΔG values for each protein and their difference in melting temperature estimated from thermal proteomic profiling (Spearman’s ρ = -0.05, p = 0.17). Similarly, predicted ΔΔG values for Guk1 (0.03) and Aha1 (0.17) were not suggestive of their measured melting temperatures.

## Discussion

Species that inhabit disparate thermal environments exhibit evidence of thermal adaptation based on differences in their induction of the heat shock response and in their protein melting temperatures. Prior studies have refined these observations by identifying specific proteins that are thermal sensors and have diverged between even closely related species [1,2,5]. Our results demonstrate a proteome wide shift in thermostability between two thermally diverged mesophilic *Saccharomyces* species. We also show that changes in the cellular environment contribute to protein melting temperature divergence. However, most proteome stability differences persist even in the same cellular environment. By showing that thermostability divergence is structurally encoded for Guk1 and Aha1 and is structurally predictable across the proteome, our results imply that divergence in thermotolerance involves massively concordant changes in protein stability that may constrain the rate of adaptation to high temperatures.

### Coordinated evolution of protein melting temperatures

Most thermophilic and mesophilic organisms have been separated for billions of years and exhibit large differences in protein melting temperatures throughout their proteomes [6,35,38,48]. We estimate that nearly 85% of proteins in *S. cerevisiae* are more stable than their *S. uvarum* orthologs, implying a similarly massive set of coordinated changes in protein thermostability between relatively closely related yeast species. This estimate excludes proteins with inconsistent differences between replicates but is fairly uniform across protein groups with the exception of ribosomal proteins. The melting temperature differences in the hybrid, while smaller in magnitude, are largely concordant with the parental species differences and indicate that dominant cellular factors from *S. cerevisiae* stabilize *S. uvarum* proteins in the hybrid. This cellular dominance could contribute to the hybrid’s ability to grow at high temperature (37°C) [49]. While previous studies of melting temperature divergence were not able to account for the contributions of cellular environment, our results are not surprising given that prior studies have shown that cell type, treatment and metabolic state can influence protein melting temperatures [6,15,36,37]. However, our results are also limited as they do not address in vivo stability, where the concentration of cellular factors is important. Further work will be needed to identify how cell environments diverge between species, as such changes could play an important role in the evolution of thermotolerance.

### Fitness consequences of protein stability

A central concept from comparative studies is that selection shapes the distribution of protein melting temperatures across species. However, selection on melting temperatures is most likely indirect and caused by selection on protein stability or other biophysical properties at physiological temperatures since most protein melting temperatures greatly exceed an organism’s thermal growth limits. Prior studies have shown that divergence in protein melting temperatures can result from changes in its heat capacity and/or enthalpy and is most likely driven by selection to prevent misfolding and maintain protein activity [35,50,51]. This biophysical model of protein evolution is supported by the observation that most proteins are marginally stable such that most mutations are destabilizing [20,22,52].

Consistent with the idea that selection acts to maintain levels of protein stability at the temperatures experienced by an organism, we find that amino acid differences between the two species are predicted to stabilize *S. cerevisiae* or destabilize *S. uvarum* proteins. However, we also find no correlation between protein melting temperature differences and the sum of the predicted ΔΔG across a protein. Besides measurement error, the absence of a correlation could indicate that changes in protein stability are not additive or that the temperature of aggregation (Tm) differs from the temperature of unfolding (ΔΔG) in a protein-specific way. Prior studies have shown that non-additive (epistatic) effects among protein substitutions are common and increase as protein homologs diverge [53,54].

Proteins vary in how changes in stability are related to fitness, with some proteins being more robust to changes in stability than others [53]. In the case of Guk1, we find no detectable fitness effects caused by placing the thermally sensitive *S. uvarum* allele into an *S. cerevisiae* background. One explanation is that *S. uvarum* Guk1 is less stable at physiological temperatures but protein levels are not sufficiently reduced to generate detectable fitness effects. Prior studies have shown that while null alleles of *GUK1* are recessive lethal [55], temperature sensitive alleles of *GUK1* are unstable and degraded by the proteasome at restrictive temperatures [56]. Thus, even if thermally sensitive *S. uvarum* alleles do not substantially affect fitness at high temperatures, the combined effects of numerous thermosensitive *S. uvarum* proteins could limit thermotolerance via a denaturation catastrophe [57].

### Protein stability as a mechanism of thermotolerance

How does protein stability evolve in relation to thermotolerance? Given the preponderance of destabilizing mutations, loss of stability may easily occur as a neutral process when there is no selection for thermostability via exposure to high temperatures. Thus, one possibility is that the the ancestor of the *Saccharomyces* had moderate to high thermotolerance which was then lost along with protein stability on each of the lineages leading to thermally sensitive species. Consistent with parallel loss of stability, *S. kudriavzevii* proteins aggregate at a lower temperature in vivo than *S. cerevisiae* proteins [5]. However, the results of this study show an adaptive role in tuning a species’ heat shock response. Another, non-exclusive evolutionary scenario is gain of thermotolerance along the lineage leading to *S. cerevisiae* (Figure 1)[58]. In the context of increasing thermotolerance, thermostability may impose a significant constraint on the rate of adaptation: a mutation that enhances the stability of one protein may not have much impact on thermotolerance without concordant changes in the rest of the proteome. Thus, stabilization of the entire proteome may be constrained to incrementally occur through many changes of small effect. However, changes in the cell environment, e.g. due to thermoprotectants like trehalose [59] or protein chaperones like Aha1 [60], provide one mechanism that could alleviate the constraints of coordinated changes throughout the proteome by generating a temporary buffer that facilitates the evolution of higher thermotolerance.

## Methods

### Species’ thermotolerance

We measured thermal tolerance by the final density of cultures grown across a range of temperatures for representative strains of seven species (S2 File, A). Overnight rich medium (YPD: 1% yeast extract, 2% peptone and 2% dextrose) cultures were grown to mid-log phase and diluted 1:100 (vol:vol) in fresh medium. Cultures (100 µl) were grown in PCR tubes and incubated in a gradient thermocycler across a range of temperatures for 24 hours. For each gradient, temperatures were empirically measured to avoid slight inaccuracies in the machine estimated gradient. Final concentrations were measured by OD600 in flat bottom plates. The temperature at which growth was inhibited by 50% along with confidence intervals was estimated for each species by fitting a five parameter least squares regression using the nplr package (v0.1-7) in R [61].

### Thermal Proteome Profiling

#### Lysate preparation

We characterized proteome thermostabilities of two replicates of the haploid parental strains, *S. cerevisiae* (YJF155) and *S. uvarum* (YJF1449), and one replicate of the interspecies hybrid (YJF4680: YJF155 x YJF1449, S2 File, A). Strains were grown overnight at 25°C in YPD, resuspended in fresh medium, and harvested during log phase after 4 hours of growth. Cells were combined in equal portions with cells from the overnight culture to capture proteins expressed in both log and stationary phase cells. Cells were lysed by bead beating in lysis buffer (20 mM sodium phosphate, 50 mM NaCl, pH 7.4, 1 EDTA-free protease inhibitor tablet from ThermoFisher). The lysate was sonicated to shear any DNA present and soluble protein was collected from the supernatant after centrifugation at 18,000 g for 20 minutes at 4°C.

#### Temperature treatments

Protein lysates were quantified using the bicinchoninic assay (Pierce BCA) and diluted to 2 µg/µl in lysis buffer. Protein extracts (50 µg aliquots) were incubated at temperatures ranging from 30°C to 72°C for 3 minutes in a thermocycler (S2 file, B). We used 10 temperatures for the parental species (TMT 10plex) and 16 different temperatures for the hybrid (TMT 16plex). BSA (1 µg) was added as a standard to each tube and samples were centrifuged at 18,000 g for 20 minutes at 4°C to remove aggregated proteins.

#### Desalting and TMT tagging

The supernatant containing the soluble fraction was applied to Ultracel 10K filters (Milipore Sigma) and washed with urea buffer (8 M urea, 100 mM triethylammonium bicarbonate (TEAB, pH 8.5)) by centrifuging at 14,000 g for 15 minutes. The samples were subsequently washed with TCEP buffer (5 mM TCEP (CAS #51805-45-9), 8 M urea, 100 mM TEAB), IAA buffer (50 mM iodoacetamide (IAA), 8 M urea, 100 mM TEAB) and 100 mM TEAB. Next, samples were digested with 100 µL of trypsin solution (Thermo Fisher #90058 - lyophilized powder diluted 1:50 in 100mM TEAB) overnight at 37°C. The digest was collected by centrifugation, dried down usign a SpeedVac and frozen at -80°C.

Samples were resuspended in 100 µL of 0.1% trifluoroacetic acid (TFA) in LC-MS grade water. We then activated stage tips by two washes with 100 µl of 50/50 acetonitrile (ACN) and 0.1% TFA, and two washes with 0.1% TFA. We then applied the resuspended samples to the stage tips and spun them down. Samples were washed twice with 100 µl of 0.1% TFA. All spin steps were carried out at 2200 g for 4 minutes. Next, we eluted with 100 µl of 50/50 ACN and 0.1% TFA. We again dried down and froze the samples at -80°C.

Samples were resuspended in 125 µl of 100 mM TEAB. 25 µl of the sample was labeled with 20 µl of isobaric Tandem Mass Tags (TMT, Thermo Fisher) and resuspended in ACN. Parental replicates 1 and 2 were tagged with a randomized TMT10plex, whereas hybrid samples were tagged with a randomized TMT16plex. Samples were quenched with 5 µl of 5% hydroxylamine for 15 minutes after incubating at room temperature for 1 hour. The 10 (or 16 for hybrid) samples were combined, dried down and frozen.

#### Fractionation

The combined sample was chemically fractionated into 8 fractions using a stage tip activated with two washes of 50 µl of ACN and two washes of 50 µl of 100 mM ammonium formate (AF). The sample was resuspended in 50 µL of 100 mM AF and applied to the activated stage tip. For the parental replicates, 8 elution buffers (E1-E8) and a wash buffer were prepared using varying ratios of acetonitrile and 100 mM ammonium formate (10% ACN to 50% ACN). For the hybrid, we used 16 elution buffers (E1-E16) ranging from 2% ACN to 50% ACN (S2 File, C). After adding the wash buffer to the stage tip, we sequentially elute with 50 µl of the elution buffers (E1-E8 or E1-E16), collecting each flow-through in a new tube. For all these steps, we spun samples at 2200 g for 2 minutes. For the hybrid, we combined fractions E1 and E9, E2 and E10, etc., to end up with 8 fractions. The 8 eluted fractions were frozen, dried down in a SpeedVac, and resuspended in 0.1% TFA and processed at the University of Rochester Mass spectrometry core for LC/MS-MS.

#### LC/MS-MS

Peptides were injected onto a homemade 30 cm C18 column with 1.8 um beads (Sepax), with an Easy nLC-1200 HPLC (Thermo Fisher), connected to a Fusion Lumos Tribrid mass spectrometer (Thermo Fisher). Solvent A was 0.1% formic acid in water, while solvent B was 0.1% formic acid in 80% acetonitrile. Ions were introduced to the mass spectrometer using a Nanospray Flex source operating at 2 kV. The gradient began at 3% B and held for 2 minutes, increased to 10% B over 7 minutes, increased to 38% B over 94 minutes, then ramped up to 90% B in 5 minutes and was held for 3 minutes, before returning to starting conditions in 2 minutes and re-equilibrating for 7 minutes, for a total run time of 120 minutes. The Fusion Lumos was operated in data-dependent mode, with both MS1 and MS2 scans acquired in the Orbitrap. The cycle time was set to 2 seconds. Monoisotopic Precursor Selection (MIPS) was set to Peptide. The full scan was collected over a range of 400-1500 m/z, with a resolution of 120,000 at m/z of 200, an AGC target of 4e5, and a maximum injection time of 50 ms. Peptides with a charge state between 2-5 were picked for fragmentation. Precursor ions were fragmented by higher-energy collisional dissociation using a collision energy of 38% and an isolation width of 1.0 m/z. MS2 scans were collected with a resolution of 50,000, a maximum injection time of 105 ms, and an AGC setting of 1e5. Dynamic exclusion was set to 45 seconds.

### Data Analysis

#### Peptide processing

MaxQuant (v1.6.17.0) was used to identify peptides from the LC/MS-MS data and map them to reference proteins of *S. cerevisiae* and *S. uvarum* nuclear [62] and mitochondrial genomes (NC_027264 and NC_031512). We used the default parameters for modifications, setting the group specific parameter type to Reporter ions MS2 and setting digestion type to Trypsin (see TableS10 for parameters). The hybrid LC/MS-MS data was separately mapped to *S. cerevisiae* and *S. uvarum* and a modified MaxQuant xml file was used to handle the TMT16plex data [63]. For the hybrid dataset, peptides that mapped to both *S. cerevisiae* and the *S. uvarum* proteome were removed (43% of detected peptides). Peptide abundance was extracted from MaxQuant evidence files and used as input for MSTherm (v0.4.7), an R package used to analyze TPP data [64]. To compare proteins between species we used orthologs previously determined by homology and synteny [62]. From 5261 orthogroups, we removed 24 due to either highly similar paralogs, large differences in protein length or small regions of homology compared to overall protein length. *S. cerevisiae* gene names from the Saccharomyces Genome Database (SGD) were used for all orthologous protein pairs.

#### Measuring protein melting temperatures

All three datasets were standardized (to BSA spike in) and normalized using MSTherm. Proteins with three or more peptide spectra matches (PSM) were retained. To compare the three datasets, the melting temperatures of the hybrid and second parental replicate were normalized to the first parental replicate. This was accomplished by first estimating Tm values for each protein in the datasets separately, then fitting a linear regression between the Tm values in the first parental replicate and the replicate to be normalized. Importantly, each of the three sets included both *S. cerevisiae* and *S. uvarum* melting temperature values, resulting in equivalent adjustments of each. The regression was then used to adjust the hybrid and second parental set of temperature values. Finally, normalized melting temperature estimates were obtained for each dataset using the adjusted temperature values. The temperature vectors pre and post normalization are listed in S2 File, B. All melting temperatures were obtained using the four parameter logistic function in MSTherm.

#### Filtering criteria

We removed proteins with missing or low quality melting temperatures after normalization. First, we removed proteins that didn’t have orthologs. Second, we removed proteins with r^2^ < 0.8 for the logistic fit of the melting temperature curve. Third, we removed proteins that only had data in one species within a given replicate. For the two parental replicates this resulted in 827 protein pairs, and for all three (parents + hybrid) this resulted in 344 orthologous protein pairs. The smaller size of the hybrid dataset was caused by fewer proteins with at least three species specific PSMs.

MSTherm estimates 95% confidence intervals for proteins with at least eight PSMs using bootstrap resampling. There were 462 proteins with confidence intervals in both parental replicates, of which 206 had non-overlapping confidence intervals between species in both replicates. Only 129 proteins had confidence intervals in the hybrid and the intersection of proteins with non-overlapping confidence intervals in all three datasets was 34.

#### Protein annotations

Proteins were classified based on whether they are essential, membrane associated, mitochondrial, glycosylated, ribosomal or part of a protein complex. A manually curated set of 1900 proteins in 616 complexes in yeast was obtained from the Complex Portal [65]. Out of 827 proteins we examined, 310 were in one or more of these complexes. SGD was used to identify membrane associated proteins (149), mitochondrial proteins (127), subunits of the ribosome (62), and to classify proteins as essential (246), non-essential (527) or undetermined (54). Using data from Neubert et al. [66] 98 of the 827 proteins were annotated as glycosylated. Aggregation prone proteins (super aggregators) were obtained from Wallace et al. [67]. Protein abundance was obtained from [68] and the nonsynonymous substitution rate was used as a measure of protein divergence [43].

### Protein purification

#### Cloning

We cloned *S. cerevisiae* and *S. uvarum* alleles of *GUK1* and *AHA1* into a protein overexpression vector, AVA421 [69] digested with *PmeI* and *NruI*, using NEBuilder HiFi DNA Assembly Cloning kit. This vector has an IPTG inducible promoter with an N terminal 6xHis tag. Assembled plasmids were transformed into DH5-alpha *E. coli* (NEB) and sequenced. Confirmed plasmids were then transformed into BL21 (DE3) *E. coli* (NEB) for overexpression and purification.

#### Cell harvest and protein purification

Three liters of LB supplemented with 100 μg/mL of ampicillin was inoculated with 7 mL/L of overnight culture of BL21 (DE3) *E. coli* carrying an AVA421 plasmid with *GUK1* or *AHA1* from *S. cerevisiae* or *S. uvarum,* and grown at 30°C until it reached an OD600 ∼0.5. Next, protein expression was induced using 1 mM IPTG and the cultures were grown overnight at 18°C. The next day, cells were spun down, washed with water and quickly frozen on dry ice. The pellet was resuspended in sonication buffer (20 mM HEPES 7.5, 1 M NaCl, 5% glycerol, 2 mM BME, 2 μg/mL Leupeptin, 1 μg/mL Pepstatin and 1 mM Pefabloc) and sonicated to lyse cells for 10 seconds for a total of 8 cycles with 1 minute of rest in between on ice.

Sonicated cells were spun down and the supernatant was diluted with an equal volume of buffer without salt (20 mM HEPES pH 7.5, 5% glycerol, 2 mM BME). The supernatant was applied to 2 mL slurry of TALON Metal Affinity Resin (Takara Bio #635504) and nutated for 1 hour to allow TALON to bind the His tagged protein. We then removed the supernatant and washed the resin four times with 25 mL 0.5 M NaCl wash buffer (20 mM HEPES pH 7.5, 5% glycerol, 0.5 M NaCl, 2 mM BME), nutating at 4°C for 10 minutes. The sample was washed with 25 mL of E1 buffer (5 mM imidazole pH 7.7 in 0.5 M wash buffer) followed by 25 mL of E2 buffer (10 mM imidazole pH 7.7 in 0.5 M wash buffer) and applied to the TALON on a column. We eluted the protein by applying E3 buffer (250 mM imidazole pH 7.7 in 0.5 M wash buffer) to the column.

Protein concentration was quantified using the Bradford assay and 55 ul of Genscript 3C His protease was added for every 10 mg of protein to cleave the 6x His tag from the protein. The sample was dialyzed into 200 mM NaCl, 20 mM HEPES pH 7.5, 5% glycerol, 2 mM BME overnight. The next day, protein was harvested, diluted with an equal volume of 0.8 M NaCl, 20 mM HEPES pH 7.5, 5% glycerol, 2 mM BME and applied to a 1.5 mL slurry of TALON. We nutated the sample for 1 hour at 4°C and vacuum filtered the sample (0.2μm). The sample was then dialyzed overnight into storage buffer (20 mM HEPES pH 7.5, 0.5 mM BME, 10 mM NaCl). The following day, protein was harvested and frozen using liquid nitrogen for long term storage at -80°C.

#### Circular Dichroism spectroscopy

We used circular dichroism (CD) spectroscopy to estimate the melting temperatures of the purified proteins. Guk1 proteins were diluted to a final concentration of 0.25 mg/mL in buffer (10 mM HEPES pH 7.5, 0.25 mM BME and 5 mM NaCl). We used a quartz cuvette (1 mm path length) and the buffer with no protein as a blank in a JASCO J-1100 Circular Dichroism spectrometer. We first performed a wavelength scan from 200 nm to 260 nm with 1 nm bandwidth and 0.1 nm step size to characterize the CD spectra of Guk1. Thermal melts were performed by tracking ellipticity at 222 nm for Guk1 from 4°C to 74°C with 1°C/min increments with a 0.2°C interval. Aggregation was irreversible when we reduced the temperature from 74°C to 4°C. Wavelength scans were also performed at seven temperatures during the thermal melt (Figure S4). Three replicates of thermal melts were done for each Guk1 ortholog.

Aha1 was not stable in our buffer and crashed out when we thawed the protein. To avoid this interfering with CD, we spun the protein solution at 4°C to remove insoluble protein and carried out thermal melts at 220 nm from 4°C to 74°C with 1°C/min increments with a 0.2°C interval. The wavelength scans were carried out from 210 nm to 260 nm with 1 nm bandwidth and 0.1 nm step size. Aggregation of Aha1 was also irreversible. Only one thermal melt was carried out for each of the two Aha1 orthologs.

We calculated mean residue molar ellipticity, by dividing the measured ellipticity (θ) by cuvette path length (l), protein concentration (c) and number of residues in the protein (N).

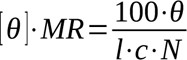

Next, we used the nls2 (v0.3-4) R package to fit a non-linear regression model to [θ]_MR_ and estimated the melting temperature. For Guk1, we calculated the average [θ]_MR_ across the three replicates and then fit a model for a two state transition from folded to unfolded states, allowing for linear changes in ellipticity pre and post transition (Model 2). For Aha1, we fit a model for a two state transition from folded to unfolded states to the calculated [θ]_MR_ (Model 1). Both models are from supplementary equations in Greenfield [70]. The estimated model parameters for all four proteins are in Table S9.

### Enzymatic activity measurements

#### Crude protein extraction and temperature treatments

*S. cerevisiae* (YJF155) and *S. uvarum* (YJF1449) cultures were grown in YPD to either mid-log (OD600 ∼ 0.6-0.8) or stationary phase (overnight) at 25°C shaking at 300 rpm. Cells were washed twice with sterile water and the pellets stored at -80°C until use. Frozen pellets were thawed and resuspended in 0.5 mL assay buffer (see assay descriptions) and transferred to Protein Lobind tubes (Eppendorf, Hamburg) containing 0.25 mL 0.5 mm glass beads. The lysates were pulverized in a Mini Bead-beater-96 (Biospec Products, Bartlesville, OK) over 5 cycles consisting of 90 seconds on ice followed by 30 seconds of homogenization. Lysates were clarified in a 4°C centrifuge for 20 minutes at 18,000 g and the protein concentration of the supernatants was determined by Pierce BCA assay (Thermo Fisher Scientific, Waltham, MA) using the assay buffers to dilute the samples and bovine serum albumin standards. Clarified samples were diluted to protein concentrations optimized for each assay and 100 uL aliquots were transferred to PCR tubes for heat treatments ranging from 25°C to 80°C performed in thermocyclers (25°C for 3 min, 3 min temperature treatment, 25°C for 3 min). Heated samples were transferred to Lobind tubes and clarified once again before loading into plates for enzymatic activity assays.

#### Malate Dehydrogenase Activity Assays

Malate dehydrogenase activity of soluble crude protein extracts was assessed using an assay kit (Sigma-Aldrich, St. Louis, MO). Clarified cell lysates were diluted to 8 µg/mL prior to heat treatment. Reaction mixes with or without malate were combined with heat treated lysates in 96-well plates and the absorbance at 450 nm was recorded every 2 minutes for an hour in a plate reader heated to 37°C.

#### Glutathione Reductase Activity Assays

Glutathione reductase activity of soluble crude protein extracts harvested from stationary cultures was assessed using an assay kit (Sigma-Aldrich, St. Louis, MO) modified for use in a 96-well plate format. Clarified cell lysates were diluted to 500 µg/mL in the included assay dilution buffer prior to heat treatment. Assay reaction buffer (100 mM potassium phosphate buffered to pH 7.5, 1 mM EDTA, 150 uM reduced β-NADPH, 1 mM DTNB) with or without 1.3 mM oxidized glutathione was added to diluted samples in a 4:1 (vol:vol) ratio in 96 well plates. The absorbance at 412 nm was recorded every 10 seconds for 15 minutes in a plate reader.

#### Guanylate kinase activity assays

Guanylate kinase activity of purified Guk1 from *S. cerevisiae* and *S. uvarum* was measured after exposure to various temperatures. Frozen proteins were resuspended in TPP buffer (20 mM sodium phosphate buffer, 50 mM sodium chloride, EDTA-free protease inhibitors, pH 7.4) and the concentration was determined by Pierce BCA assay. Protein concentrations were adjusted to 5 µg/mL with TPP buffer and subjected to heat treatments in a thermocycler as described above. Residual guanylate kinase activity was determined using the assay described by Lecoq et al [71] with adaptations made for a multi-well plate format. Assay buffer (100 mM Tris-HCl pH 7.5,100 mM KCl, 10 mM MgCl_2_, 300 uM reduced β-NADH, 600 uM PEP, 2 mM ATP, 35 units pyruvate kinase from rabbit muscle, 80 units lactate dehydrogenase from rabbit muscle, and 880 uM GMP) was added to heat-treated samples in a 10:1 vol:vol ratio in 96 well plates. The absorbance at 340 nm was recorded every 10 seconds for 15 minutes in a plate reader heated to 30°C. Guk1 kinetics were determined by combining non-heat treated samples with the assay buffer without GMP in a 1:9 vol:vol ratio in a 96-well plate, pre-incubating the plate at temperature (30°C or 37°C) for 5 minutes, followed by initiating the assay with an equal sample volume of GMP for final concentrations ranging from 5 uM to 10 mM and measuring the absorbance at 340 nm every 10 seconds for 15 minutes in a plate reader heated to the pre-incubation temperature. We were unable to find conditions where Guk1 activity could be measured at high temperatures. Magnesium was previously identified to enhance the thermal stability of various proteins [72] in addition to increasing thermotolerance [73,74] and we observed that it enhanced the residual enzymatic activity of Guk1 after exposure to higher temperatures when added to TPP buffer. However, magnesium is required by pyruvate kinase [75] which produces pyruvate in proportion to the GDP produced by Guk1 and is used to generate the observable 340 nm absorbance change as β-NADH is converted to β-NAD by lactic dehydrogenase. We found that magnesium chloride concentrations above 2.5 mM caused measurable Guk1 thermoprotection while concentrations below 2.5 mM rendered pyruvate kinase to become the rate-limiting reaction in the assay.

#### Enzymatic activity assay analyses

Enzymatic activity in a reaction was defined as the maximum change observed across 6-12 consecutive recorded time points. Activity rates for assays performed with crude protein extracts were corrected for intrinsic or off-target activity by subtracting maximum changes determined from substrate blank reactions performed with the same heat-treated sample and normalized to the rate observed for samples heat challenged at 25°C. The inhibitory temperature at which 50% of the maximum observed activity remained was calculated using 5 point logistic regression via the nplr R package [61]. Statistical significance was tested using Students’ 2-tailed t-test.

### *GUK1* allele swap and phenotyping

#### Allele swap

We generated an allele replacement of *S. cerevisiae GUK1* with the *S. uvarum GUK1* in an *S. cerevisiae* background (YJF4559 - Mat alpha hoΔ::dsdAMX4 ura3Δ derived from YPS163) under the native *S. cerevisiae* promoter. Since *GUK1* is essential, we used CRISPR to cut within *S. cerevisiae GUK1* (onw37 and onw38 in S2 File, D, see Laughery et. al [76] for cloning details) and provided a repair template consisting of *S. uvarum GUK1* (564 bp) flanked by ∼ 200 bp of *S. cerevisiae* upstream and downstream sequence to increase the chance of recombination occurring outside the *GUK1* coding sequence. This repair template was generated using NEB HiFi DNA assembly on pRS316 digested with BamHI and HindIII, three PCR purified fragments of ∼200bp sequence upstream of *S. cerevisiae GUK1*, *S. uvarum GUK1* coding sequence and ∼200bp sequence downstream of *S. cerevisiae GUK1*, and then amplified using external primers. We then screened transformants using allele specific PCR [77] and Sanger sequenced Guk1 and flanking regions (primers listed in S2 File, D). Out of 14 screened transformants, 9 were total gene replacements and one was a chimera (63.8% N terminal nucleotide sequence from *S. uvarum GUK1*). We used the same strategy to generate an allele replacement in *S. uvarum* (YJF4581) and obtained four full length allele replacements and one chimera (83% N terminal sequence from *S. cerevisiae GUK1*) which were all confirmed by Sanger sequencing.

#### Phenotyping

*S. cerevisiae* (YJF4559), *S. cerevisiae* with *S. uvarum GUK1* (YJF5540) and *S. cerevisiae* with chimeric *GUK1* (YJF5548) were phenotyped at multiple temperatures 30°C, 37°C, 39°C and 41°C as well as at room temperature subsequent to a heat shock. For the constant temperature assays, strains were arrayed in a 96 colony format and a Singer Rotor HDA (Singer Instruments, Somerset, England) was used to pin the strains to agar plates made with synthetic complete media (CM), minimal media (MM) supplemented with uracil (50 mg/L), and minimal media supplemented with uracil and mycophenolic acid (MPA, 100 µg/mL), an inosine monophosphate dehydrogenase and guanosine biosynthesis inhibitor [78]. Plates were incubated at four temperatures for three days and colony size was measured using images obtained from a Singer Phenobooth (Singer Instruments). For the heat shock assays, the strains were arrayed on CM agar plates, incubated at 50°C for 20, 40, 60, 100, 160 and 300 minutes and then shifted to room temperature for an additional two days. Colonies on the edge of the plate as well as one row/column in were removed from the analysis to avoid edge effects [79]. Colony size differences were tested using an analysis of variance with a linear model: size ∼ strain*temperature for the constant temperature and size ∼ strain separately for each heat shock duration. For allele replacements in *S. uvarum*, temperature-dependent growth differences were measured by spot dilutions on minimal medium plates supplemented with uracil at 25°C, 30°C, 31°C, 32°C, 33°C and 34°C.

### Protein stability predictions

Protein structure predictions were generated using ColabFold (v1.5.2, https://github.com/YoshitakaMo/localcolabfold)[45]. Protein sequences were obtained from complete and annotated genomes of *S. cerevisiae* (DBVPG6765) and *S. uvarum* (YJF1449 = CBS7001)[80]. ColabFold was run without templates and with amber for structure refinement (relaxation/energy minimization). We obtained structures for both species orthologs for 5,097 out of 5,121 proteins.

Predicted changes in stability (ΔΔG) were obtained for each *S. uvarum* amino acid substituted into the *S. cerevisiae* protein and vice versa. Proteins were aligned using Muscle (v3.8.31)[81] and amino acid substitutions were extracted from the alignments. We used both sequence and structure based predictions from DDGun (https://github.com/biofold/ddgun)[82] and ACDC-NN (v0.0.17, https://github.com/compbiomed-unito/acdc-nn)[46,83] since they have both been shown to have good antisymmetric properties in that forward and reverse changes are opposites of each other: ΔΔG(X to Y) = -ΔΔG(Y to X)[84]. However, substitutions between species’ alleles may deviate from antisymmetry due to other amino acid differences between the two alleles. The correlation between the forward (*S. uvarum* amino acids into *S. cerevisiae* proteins) and reverse (opposite) changes was close to negative one (S2 File, E). We thus used the average of the forward and negative of the reverse change so that a negative ΔΔG indicates the *S. uvarum* amino acid is less stable than that of *S. cerevisiae*.

To control for uncertainty in predicted structures, we removed proteins with lower confidence structures and proteins with structures that differed from those originally produced by AlphaFold2 [44]. We used the per-residue confidence metric (pLDDT) from AlphaFold2 to remove 1,435 proteins with an average pLDDT across both species of less than 70. We used the lDDT metric from OpenStructure (v2.7.0)[85] to compare our *S. cerevisiae* structures with those in the AlphaFold Database [86] and removed 258 proteins with lDDT scores of less than 0.70. We used the sum of the ΔΔG predictions for each protein as a measure of protein stability differences between species. In nearly all cases, proteins with higher confidence structures showed larger stability differences between species than proteins with lower confidence structures (Table S7). To determine whether stability differences are associated with sites within regions of low or high confidence structures we examined the sum of the ΔΔG predictions across pLDDT categories using the filtered set of 3,646 proteins. For both sequence and structure based methods, the majority of the stability differences came from high confidence (pLDDT > 80) sites (Table S8).

## Data availability

Raw data and data underlying the figures is available at OSF: https://osf.io/98c6n/. Raw proteomic data and the evidence files are available through the ProteomeXchange Consortium via the PRIDE repository with the identifier PDX058629.

## Acknowledgments

Experimental data was collected with facilities and support of the University of Rochester Mass Spectrometry Resource Laboratory and the Structural Biology and Biophysics Facility. We would like to thank Emery Longan, Ethan Walker, Ruiyue Tan and other members of the Ghaemmaghami lab for discussion and feedback. We also thank Eric Hilpot for experimental advice on protein purification and Jermaine Jenkins for experimental advice on CD spectroscopy.

---

**Figure S1:**
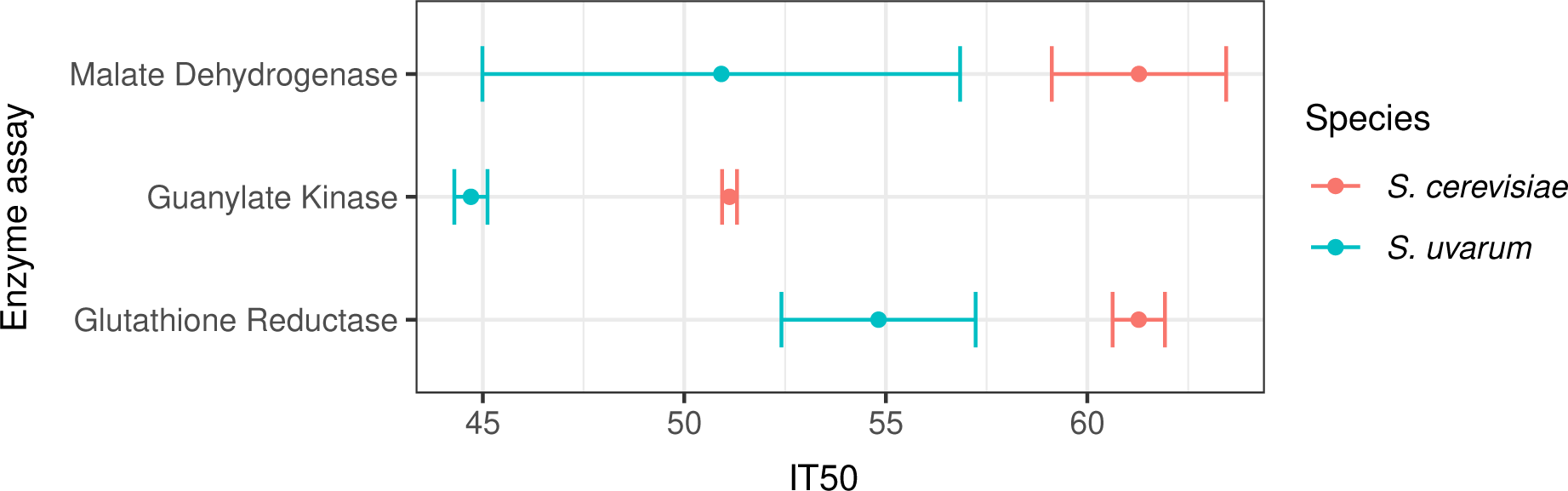
Thermal sensitivity of enzyme activity. *S. cerevisiae* and *S. uvarum* enzyme activity was measured from temperature treated lysates (malate dehydrogenase and glutathione reductase) or purified protein (guanylate kinase). The temperature at which activity was reduced 50% (IT50) is shown by the mean and its 95% confidence interval from three replicate measurements. For reference, the average of the parental species melting temperatures from the proteomics data was: 51.3 (Sc-Mdh1/3), 48.5 (Su-Mdh1/3), 50.8 (Sc-Guk1), 46.9 (Su-Guk1), 57.5 (Sc-Glr1), 49.9 (Su-Glr1).

**Figure S2:**
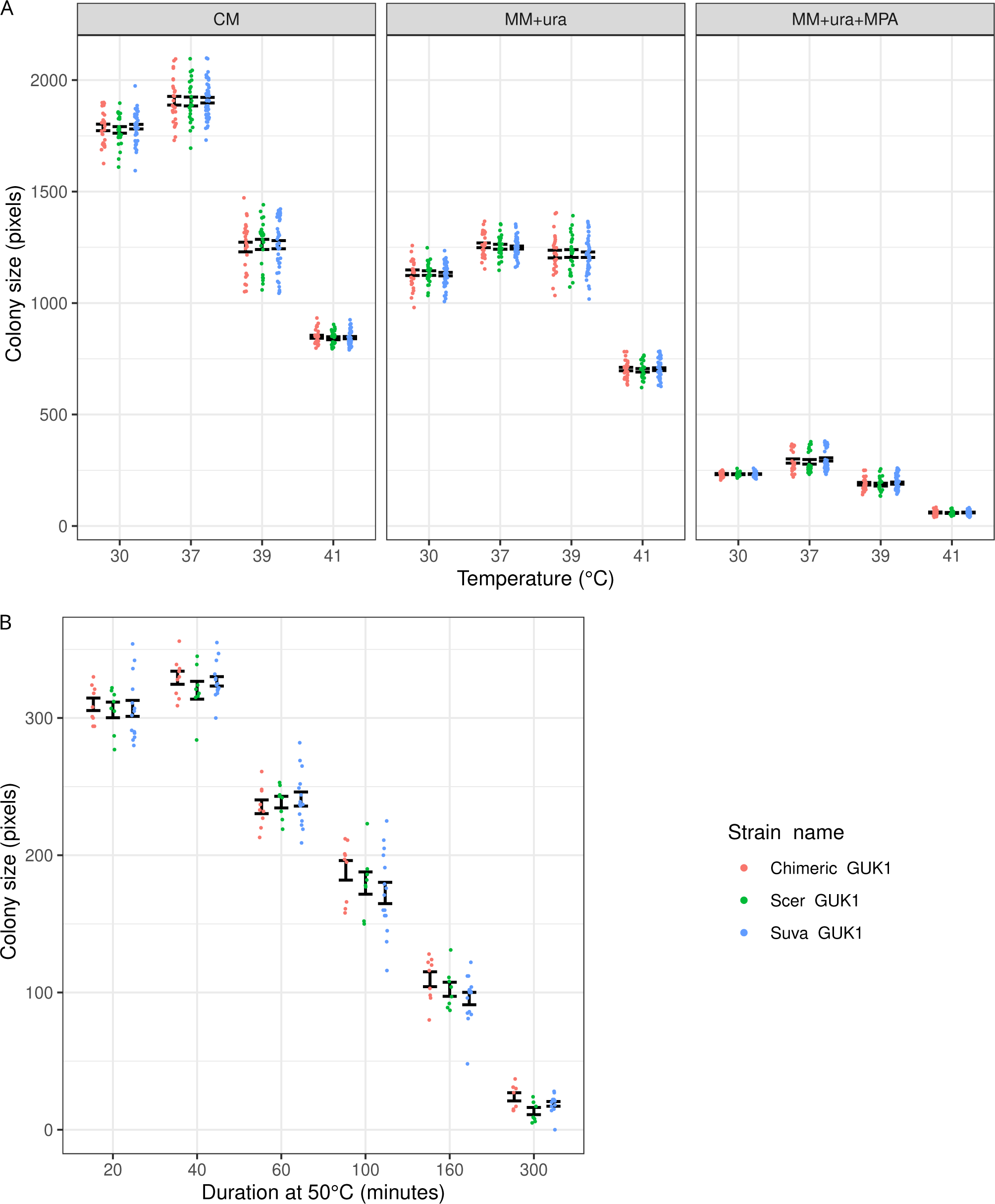
GUK1 allele-replacement phenotyping. (A) Colony size after three days of growth at different temperatures. Three panels show complete medium (CM), minimal media (MM) and minimal media with mycophenolic acid (MPA), an inhibitor of IMP dehydrogenase and synthesis of the guanosine monophosphate precursor XMP. (B) Colony size after two days of growth subsequent to a 50°C heat shock for different durations. Strains have an *S. cerevisiae* background with either *GUK1* from *S. uvarum* (Suva GUK1), *S. cerevisae* (Scer GUK1) or from a chimeric allele (Chimeric GUK1). Sample sizes are 8 (Scer), 9 (Chimera) and 15 (Suva) in panel A and 24 (Scer), 27 (Chimera), 45 (Suva) in panel B.

**Figure S3:**
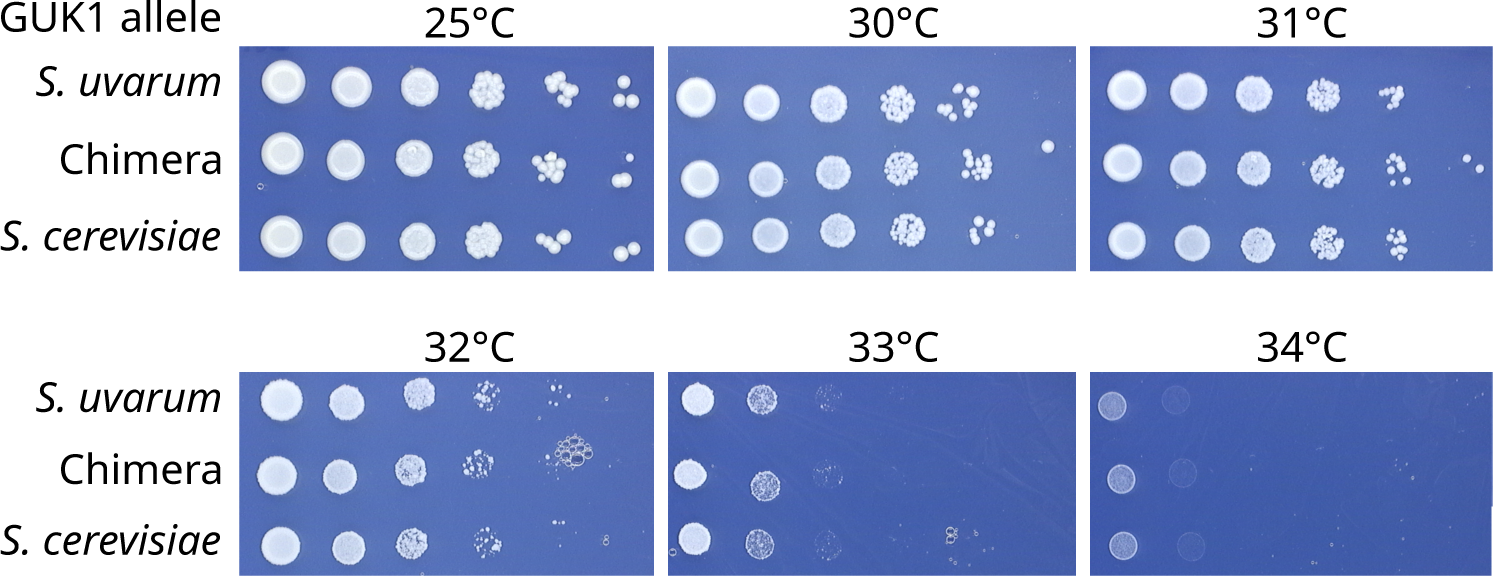
*GUK1* allele-replacement phenotyping in *S. uvarum*. *S. uvarum* strains with different *GUK1* alleles were diluted (left to right), spotted on complete medium and imaged after three days at different temperatures.

**Figure S4:**
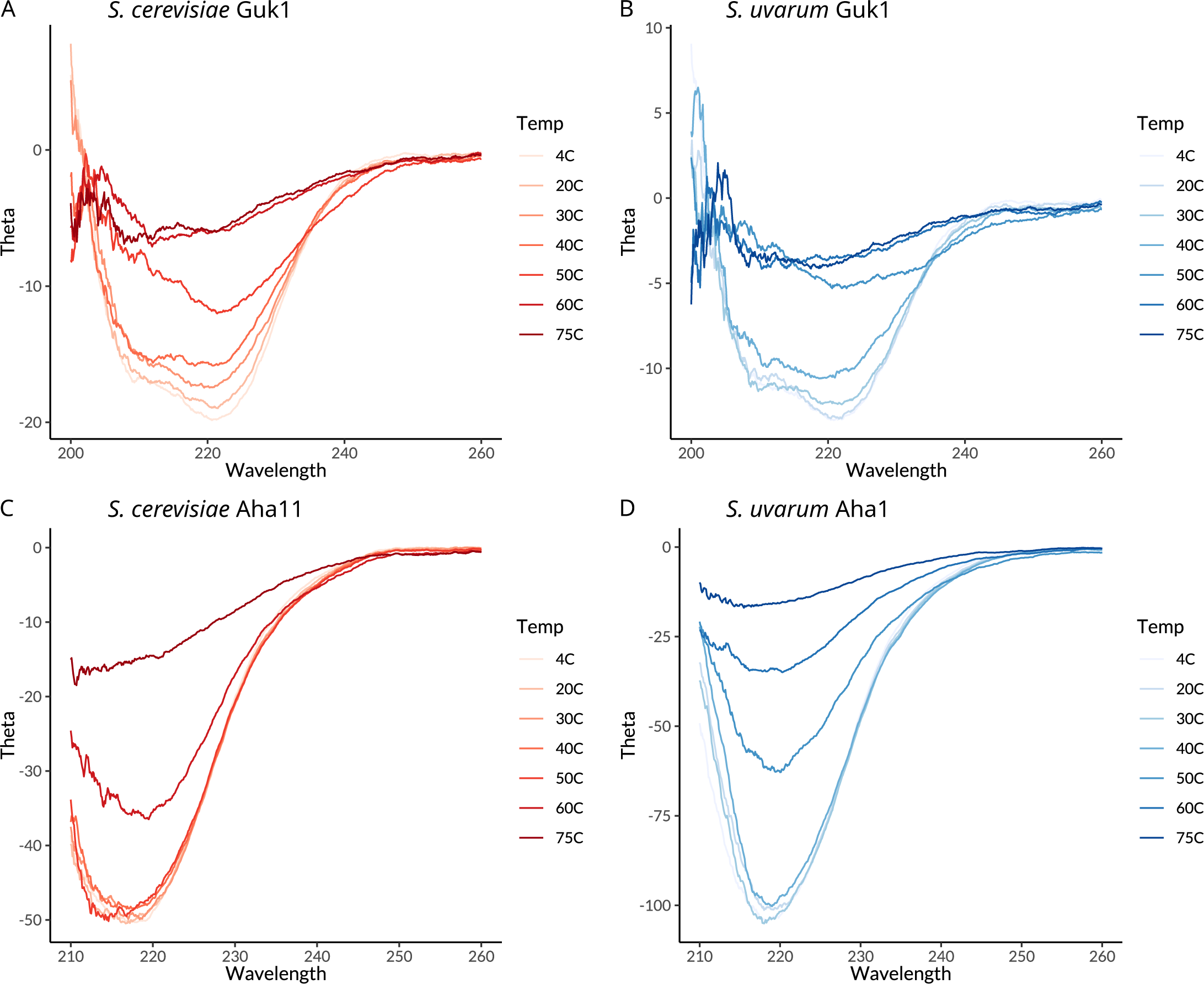
Circular dichroism wavelength scans at seven temperatures. Scans of Guk1 from *S. cerevisiae* (A) and *S. uvarum* (B), and Aha1 from *S. cerevisiae* (C) and *S. uvarum* (D) show changes in elipticity (Theta) occur at lower temperatures for the two *S. uvarum* proteins.

## Notes

https://osf.io/98c6n/

### Competing Interest Statement

The authors have declared no competing interest.

